# Does the motor cortex want the full story? The influence of sentence context on corticospinal excitability in action language processing

**DOI:** 10.1101/2020.05.21.108290

**Authors:** W Dupont, C Papaxanthis, F Lebon, C Madden-Lombardi

**Author notes:** F.L. and C. M-L. equally contributed to this work.

## Abstract

The reading of action verbs has been shown to activate motor areas, whereby sentence context may serve to either globally strengthen this activation or to selectively sharpen it. To investigate this issue, we manipulated the presence of manual actions and sentence context, assessing the level of corticospinal excitability by means of transcranial magnetic stimulation. We hypothesized that context would serve to sharpen the neural representation of the described actions in the motor cortex, reflected in context-specific modulation of corticospinal excitability.

Participants silently read manual action verbs and non-manual verbs, preceded by a full sentence (rich context) or not (minimal context). Transcranial magnetic stimulation pulses were delivered at rest or shortly after verb presentation. The coil was positioned over the cortical representation of the right first dorsal interosseous (pointer finger).

We observed a general increase of corticospinal excitability while reading both manual action and non-manual verbs in minimal context, whereas the modulation was action-specific in rich context: corticospinal excitability increased while reading manual verbs, but did not differ from baseline for non-manual verbs. These findings suggest that sentence context sharpens motor representations, activating the motor cortex when relevant and eliminating any residual motor activation when no action is present.

## 1. Introduction

Over the past couple of decades, researchers have established a clear link between language processes and perceptual-motor processes (Barsalou, 1999, 2008; Barsalou et al., 2003; Decety & Grèzes, 2006; Fischer & Zwaan, 2008; Gallese & Lakoff, 2005; Glenberg & Gallese, 2012; Jeannerod, 2008; Zwaan & Taylor, 2006). More precisely, these authors suggest that the information described by language would be treated in the brain areas that are concerned by the concept itself; if the concept concerns sight, this will lead to processing in the visual areas. The same applies to action language, such that semantic information referring to an action would engage the sensory motor system. To better understand this link, researchers have probed the involvement of the motor system during action language processing using various methodologies, such as functional Magnetic Resonance Imaging (fMRI) (Hauk et al. 2004; Pulvermüller et al. 2005; Aziz-Zadeh et al. 2006; Van Dam, Rueschemeyer, and Bekkering 2010; although for critical views see: de Zubicaray et al. 2013; Wurm and Caramazza 2019), electroencephalogram (EEG) (Kellenbach et al., 2002; Pulvermüller et al., 2001; Xu et al., 2016), behavioral measures (Andres et al., 2015; Klepp et al., 2019; Pulvermüller et al., 2001; Rabahi et al., 2012, 2013) and Transcranial Magnetic Stimulation (TMS) (Papeo et al. 2009, 2015; Labruna et al. 2011; Scorolli et al. 2012; Innocenti et al. 2014; see review: Papeo et al. 2013). Specifically, Hauk et al. (2004) using fMRI reported that silent reading of action words leads to somatotopic activation of the effector-related representation within the primary motor cortex. Papeo et al. (2009) used TMS to observe an increase in MEP amplitude for hand action verbs vs. non-action verbs 500 ms after verb presentation. These cortical activations could explain behavioral modulation during language processing, such as improvement in squat jump (Rabahi et al., 2012, 2013), and facilitation of motor response time (Andres et al., 2015; Klepp et al., 2019).

Such behavioral facilitations are present only if the linguistic content is compatible with the movement performed (Gentilucci et al. 2000; Taylor and Zwaan 2008; Dalla Volta et al. 2009; Glenberg and Gallese 2012; Zwaan et al. 2012; Van Dam and Desai 2017). Furthermore, some studies report decreases in motor activation for action verbs compared to non-action verbs at very early latencies (TMS before end of verb: Buccino et al. 2005), or fail to observe activation of the motor cortex (lexical decision tasks: Perani et al. 1999; Longe et al. 2007; motion word comprehension: Grossman et al. 2002; passive reading: Gianelli et al. 2020). While some of the discrepancies in these findings may be attributable to the timing of these studies (see early interference explanation proposed by Buccino et al. 2005), such results generally undermine the idea that action verbs systematically/automatically activate the motor cortex (see also Rüschemeyer et al. 2007; Raposo et al. 2009). Instead, this activation seems to be, as Kemmerer (2015) states, “sensitive to attentional and situational factors that we are only beginning to understand.”

In our view, context is one of these factors that plays an important role in language processing, allowing a more effective and precise understanding of the action described in the sentence. Indeed, studies using fMRI and behavioral measures demonstrate how context can modulate the degree of motor activation and performance (Beauprez et al., 2018; Gilead et al., 2013; Kemmerer, 2015; Papeo et al., 2012; Raposo et al., 2009; van Dam et al., 2010, 2012; Van Dam et al., 2010, 2012, 2014; Zwaan et al., 2010). For instance, Raposo and colleagues (2009) observed activation of motor areas for action verbs in isolation (e.g., *kick*) and in literal phrases (e.g., *kick the ball*), but not in idiomatic phrases (e.g., *kick the bucket*). Likewise, Van Dam and colleagues (2010) observed greater activation in motor-related brain areas verbs that for highly specific action verbs (to wave, to wipe) than for less specific action verbs (to greet, to clean).

Given these results, it is possible that specifying actions by increasing the sentence context may lead to greater motor cortex activation in general. Indeed, Wurm and Caramazza (2019) report a stronger decoding of action scenes relative to short sentences in frontoparietal representations, which they attribute to a difference in richness of detail: “motor-related representations might be triggered by action scenes reflecting specific aspects of the action, such as the kinematics of an action or the particular grip used on an object, whereas the motor-related representations triggered by sentences would be less specific, more variable, and less robust.” Following this logic, increasing sentence context (specifying the action) could increase motor cortex activation in general, thereby “strengthening” motor representations.

Another possibility is that greater sentence context leads to a “sharpening” of motor representations, making them more precise yet not always stronger. This idea is consistent with studies in which the context surrounding action verbs selectively engages motor representations. For instance, Taylor and Zwaan (2008) observed facilitation for rotating a knob when the direction of rotation (clockwise) matched rather than mismatched the direction described in concurrent sentences (*The gardener noticed that the water was still running. He approached the faucet which he <u>turned off</u> quickly*.). Interestingly, the motor facilitation was observed at the moment of processing the verb, and reappeared at the post-verbal adverb, only if that adverb qualified the action (“slowly” or “quickly”), and not if the adverb instead shifted focus to the agent (e.g., “obediently” or “eagerly”). This suggests that context can selectively activate motor representations, and furthermore, that these motor representations are temporally pinned to the verb, or any word that draws the contextual focus to the action itself.

In the current study, we investigated whether context leads to a general strengthening or a condition-specific sharpening of the motor representations. We used the TMS methodology to probe how context modulated activation in the motor cortex during the reading of verbs. Participants were instructed to read manual-action or non-manual verbs that appeared either in a rich sentential context (full sentence) or in a minimal context (pronoun-verb pair). TMS pulses were delivered over the pointer finger area of the left primary motor cortex at three latencies after verb presentation (200, 300, or 400 ms) to better capture individual corticospinal modulation during reading. We expect to replicate the effect of action often seen in the literature, but this effect should be modulated by sentence context. The strengthening idea would predict greater corticospinal excitability in the rich sentential context than in the minimal context for both manual and non-manual verbs. The sharpening idea, on the other hand, would predict an intensified action effect in the rich context, with an increase of corticospinal excitability for manual verbs but not for non-manual verbs.

## 2. Material and method

### 2.1. Participants

Thirty-five healthy right-handed adults (12 women; mean age = 23.79 years-old; range 18-29 years) participated in the experiment at the laboratory. Handedness was assessed by the Edinburgh inventory (Oldfield, 1971). All participants had normal or corrected-to-normal vision, without neurological, physical and cognitive pathologies. The experiment was conducted in two sessions: 1) an experimental TMS recording and 2) a reading comprehension assessment. Volunteers were first approved for participation by a medical doctor, and confirmed their participation with written consent. The Ethics Committee approved experimental protocol (CPP 2017-A00064-49) and procedures in accordance with the Declaration of Helsinki.

### 2.2. Stimuli

One hundred and forty-four French verbs were generated; half referred to *Manual Action* (e.g., “I scrub”) and half were *Non-Manual* (e.g., “I ignore”, “I kick”). These verbs were presented either in a *Rich context* that helped to specify the verb (e.g., “I see a stain on my shirt and I scrub it” or “I see a stain on my shirt and I ignore it”), or in *Minimal context* (e.g., “I scrub” or “I ignore”; see Table 1 for details). Sentences always appeared in the first-person present tense. All rich-context sentences were created such that the target verb occurred at the end of the sentence (see Figure 1). This final pronoun-verb segment was presented alone on a subsequent screen after the beginning of the sentence was presented, thus yielding an identical or very similar presentation screen to the minimal sentence version.

**Table 1.** Example of lexical stimuli used in the experiment. In grey lines are the English translation of the French verbs/sentences presented.

| Manual action |  | Non-manual |  |
| --- | --- | --- | --- |
| Minimal Context | Rich Context | Minimal context | Rich context |
| Je mélange | Les cartes sont sur la table et je les mélange | Je memorise | Les cartes sont sur la table et je les mémorise |
| *I shuffle* | *The cards are on the table and I shuffle them* | *I memorize* | *The cards are on the table and I memorize them* |
| Je frotte | Je vois une tache sur mon t-shirt et je la frotte | J'ignore | Je vois une tache sur mon t-shirt et je l'ignore |
| *I scrub* | *I see a stain on my shirt and I scrub it* | *I ignore* | *I see a stain on my shirt and I ignore it* |
| Je ferme | L'enveloppe est prête et je la ferme | Je relis | L'enveloppe est prête et je la relis |
| *I close* | *The envelope is ready and I am closing it* | *I reread* | *The envelope is ready and I reread it* |
| J'accorde | La nouvelle guitare est très belle et je l'accorde | J'écoute | La nouvelle guitare est très belle et je l'écoute |
| *I tune* | *The new guitar is very beautiful and I tune it* | *I listen* | *The new guitar is very beautiful and I listen to it* |
| Je paraphe | Après avoir lu le contrat, je le paraphe | Je contemple | Après avoir lu le contrat, je le contemple |
| I initial | After reading the contract, I initial it | I contemplate | After reading the contract, I contemplate it |

**Figure 1.**
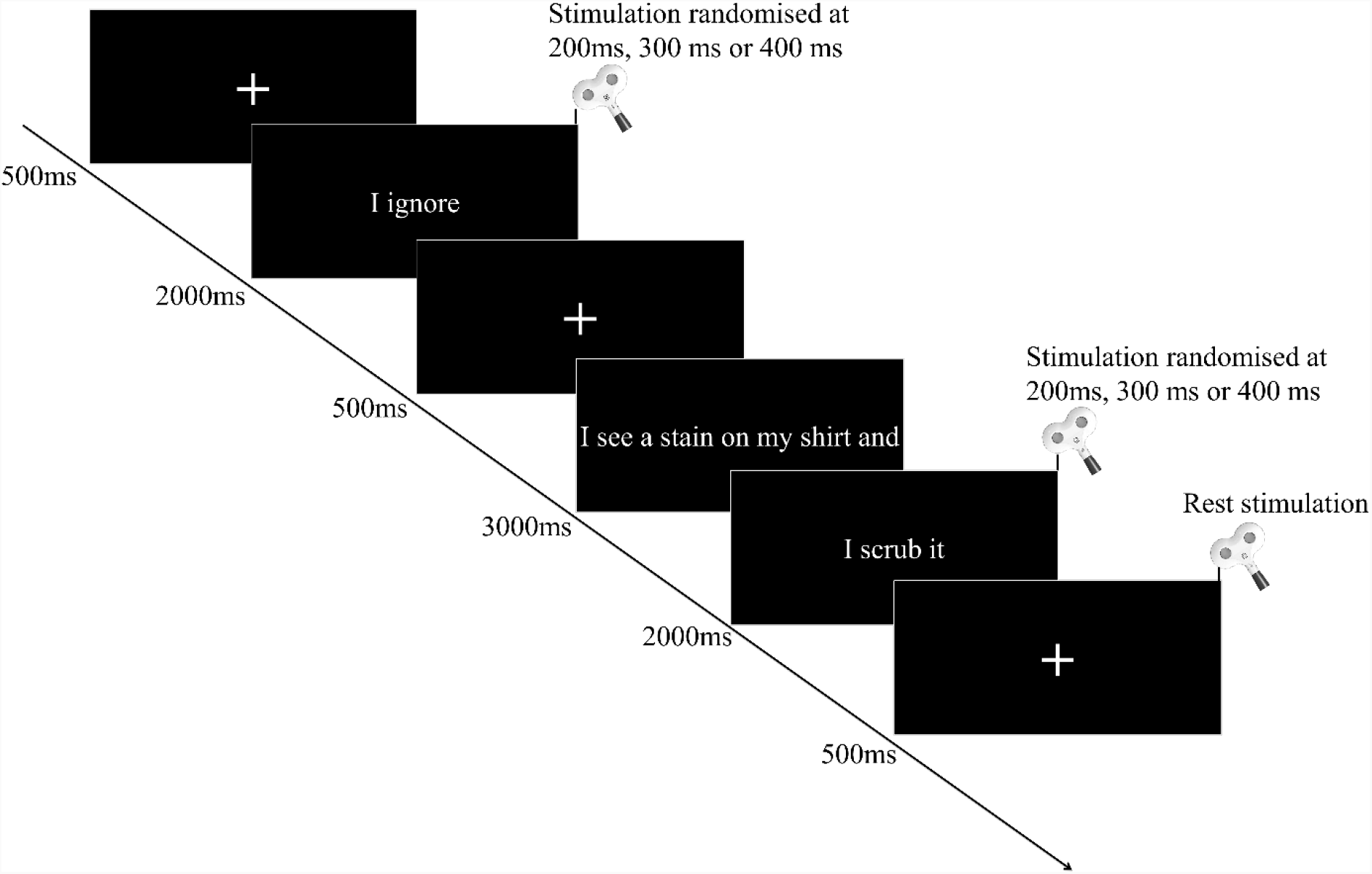
Experimental procedure. Sentences in a Minimal or Rich context were presented on a screen in front of the participant. Single-pulse TMS was triggered over the left hemisphere and motor-evoked potentials recorded in the right index finger.

A list was generated, in which 36 of the manual action verbs, and 36 of the non-manual verbs occurred within rich sentence contexts, while the other 36 manual action verbs, and 36 non-manual verbs occurred in minimal sentence context. A second list presented these same verbs in the opposite rich and minimal context versions. Each manual action verb (to scrub) was paired with a non-manual verb (to ignore), sharing the same rich-context sentence (I see a stain on my shirt and I scrub/ignore it), and each verbal stimulus only appeared once in the list. Thereby, across participants, each verb appeared in both conditions, but a given participant saw each verb and each sentence only once. For example, for the scrub/ignore pair, if a given participant read the minimal sentence “I scrub”, she/he would also read the rich context sentence “I see a stain on my shirt and I ignore it”. And another participant would read “I ignore” as well as “I see a stain on my shirt and I scrub it”. In addition, TMS latencies were counterbalanced within each list. We compared various psycholinguistic factors (written frequency, number of characters, and spelling neighbors) between manual action verbs and non-manual verbs using the Lexique.org database (New et al., 2004). Corrected t-tests yielded a significant difference only for written frequency (p=0.004; Mean ratios: 11.34 ±16.69% and 30.80±53.70% for manual action verbs and non-manual verbs, respectively), but not for number of characters (p=0.604; Mean ratios: 6.37 ±1.63% and 6.51±1.57% for manual action verbs and non-manual verbs, respectively) nor spelling neighbors (p=0.136; Mean ratios: 5.95 ±5.03% and 4.73±4.74% for manual action verbs and non-manual verbs, respectively). The frequency difference is largely due to a few high-frequency non-manual verbs such as “to read,” and this difference is in the opposite direction of the hypothesized action effect.

### 2.3. Procedure

Participants sat in an armchair. Stimuli were presented on a 19-inch LCD monitor by a home-made software, which controlled TMS triggering and synchronized physiological recordings. Throughout the recording, participants were instructed not to move, while they silently read sentences (passive reading). Corticospinal excitability was recorded at rest, as well as during the reading task at various stimulation latencies (200, 300 or 400ms after the verb onset). These latencies were chosen based on electroencephalogram studies showing 200ms as a minimal latency for semantic process in anterior regions (O Hauk et al., 2006; Pulvermüller et al., 2001). In addition, later latencies are related to changes of P300 in parietal and frontal regions (Pulvermüller et al., 2001), but also N400 in posterior regions (Beres, 2017; Kellenbach et al., 2002) when participants read action words compared to abstract words. In the present study, these different latencies were used to account for individual differences in the speed of semantic integration, which would yield variable peaks of motor cortex involvement.

Before the experimental trials, the participants performed four familiarization trials, each with a fixation cross, followed by a minimal sentence or a rich sentential context, then a black screen before the next trial. Once the participant understood this procedure, four new practice trials were presented, now including TMS pulses after the appearance of the target verb. Then, the experimental session was divided into 3 blocks, each comprised of 56 trials, yielding 168 trials total in the experiment. Among the 56 trials in each block, 8 trials were interspersed with TMS pulses only at rest (fixation cross), which served as reference stimulations and allowed comparisons across experimental conditions in that block. Therefore, participants saw 48 experimental trials in each of the three blocks, and 144 trials in total. Moreover, to avoid mental fatigue and lack of concentration, there was a two-minute break between each block, allowing participants to be as attentive as possible during experimental blocks. In each block, the number of stimulations was equally distributed between all conditions, but also between the three different stimulation latencies.

Individuals participated in a second experimental session, in which two levels of contextual reading comprehension ability were assessed. To investigate sensitivity to lower-level memory-based context, participants performed a “Remember versus Know” paradigm (Tulving, 1985),, in which they had to identify whether words were encountered in a previously read text (yes or no response), and in what context they were encountered. To assess higher-level use of context during comprehension, we used a Curriculum-Based Measurement-Maze (Parker et al., 1992), in which readers rely on inferences to select a contextually-appropriate word from three alternatives. After the first sentence, every seventh word in the passage is replaced with that correct word and two distractors. Participants must use the context to select the word that fits best with the rest of the passage.

### 2.4. TMS

Single-pulse TMS was generated from an electromagnetic stimulator Magstim 200 (Magstim Company Ltd, Whitland) and using a figure-of-eight coil (70 mm in diameter). The coil was placed over the contralateral left hemisphere to target the motor area of the First Dorsal Interosseous (FDI) muscle of the right hand. The coil rested tangential to the scalp with the handle pointing backwards and laterally at a 45° angle away from the midline. First, the mapping of the motor cortex of each participant determined the precise stimulation site (hotspot) corresponding to the location, where the MEPs amplitude of the FDI muscle was the highest and the most consistent for the same stimulation intensity. The resting motor threshold of each participant was determined as the minimal intensity of TMS necessary to induce a MEP of 50µV peak-to-peak amplitude in the right FDI muscle for 4 trials out of 8. During the experimental session, TMS intensity was set at 130% of the resting motor threshold.

### 2.5. EMG recording

The EMG signal was recorded through 10mm-diameter surface electrodes (Contrôle Graphique Médical, Brice Comte-Robert, France) placed over the FDI muscle of the right hand. Before placing the electrodes, the skin was shaved and cleaned in order to reduce noise in the EMG signal (< 20μV). The EMG signals were amplified and bandpass filtered on-line (10-1000 Hz, Biopac Systems Inc.) and digitized at 2000 Hz for off-line analysis. We also measured the root mean square of EMG signal (EMGrms), to ensure that participants were at rest in all conditions.

### 2.6. Data and statistical analysis

EMG data were extracted with Matlab (The MathWorks, Natick, Massachusetts, USA) and we measured peak-to-peak MEP amplitude. Data falling 2 SDs above or below individual means for each experimental condition were removed before analysis (4.13% of total). Then, the average MEP amplitude for each condition was normalized with reference to baseline MEP amplitude (rest). Importantly, no analysis on the timing factor was performed as we rather isolated the peak of excitability among the three stimulation times for each participant in each condition. Individuals can vary greatly in speed of decoding, as well as lexical access (Dambacher et al., 2006; Sereno et al., 1998) and semantic integration (Brown & Hagoort, 1993; Kutas & Hillyard, 1980; Van Berkum et al., 1999). This subject-specific peak method accounts for such individual variability in the semantic processing of the action over time, and allows us to tap into the action representation at the peak of the semantic representation in terms of the implication of the motor system. Finally, Grubbs test identified extreme mean values for one participant, who were excluded from the final analysis. All data analyses were performed using the software Statistica (Stat Soft, France).

Data normality was confirmed by the Shapiro-Wilk test. To assess the influence of *Action* and *Context* on corticospinal excitability, we performed a 2 by 2 repeated measures ANOVA with *Action* (Manual action vs. Non-manual) and *Context* (Minimal vs. Rich) as within-subject factors. Then, one-sample t-tests were used to compare normalized MEPs at zero for each condition to assess whether corticospinal excitability during reading changed from baseline. Finally, in order to ensure that our results were not contaminated by muscular pre-activity, we tested with Wilcoxon tests if the EMGrms before the TMS artifact was not different from zero in our experimental conditions.

To assess the influence of participants’ comprehension ability on their corticospinal excitability modulations by context, we performed Pearson correlations between the context effect (Minimal vs. Rich) and the memory-based, as well as the higher context-level comprehension scores. The data are presented as mean values (±standard error) and the alpha value was set at 0.05.

## 3. Results

The overall ANOVA revealed a main effect of *Action* (F1,33 =4.811, p=0.035, ηp²=0.127), with larger MEP ratios for manual action verbs (11.02 ±3.25%) than non-manual verbs (6.76 ±3.19%). We did not observe a main effect of *Context* (F1,33 =0.168, p=0.683, ηp²=0.005) nor an *Action* by *Context* interaction (F1,33 =0.064 p=0.801, ηp²=0.001) (Figure 2).

**Figure 2.**
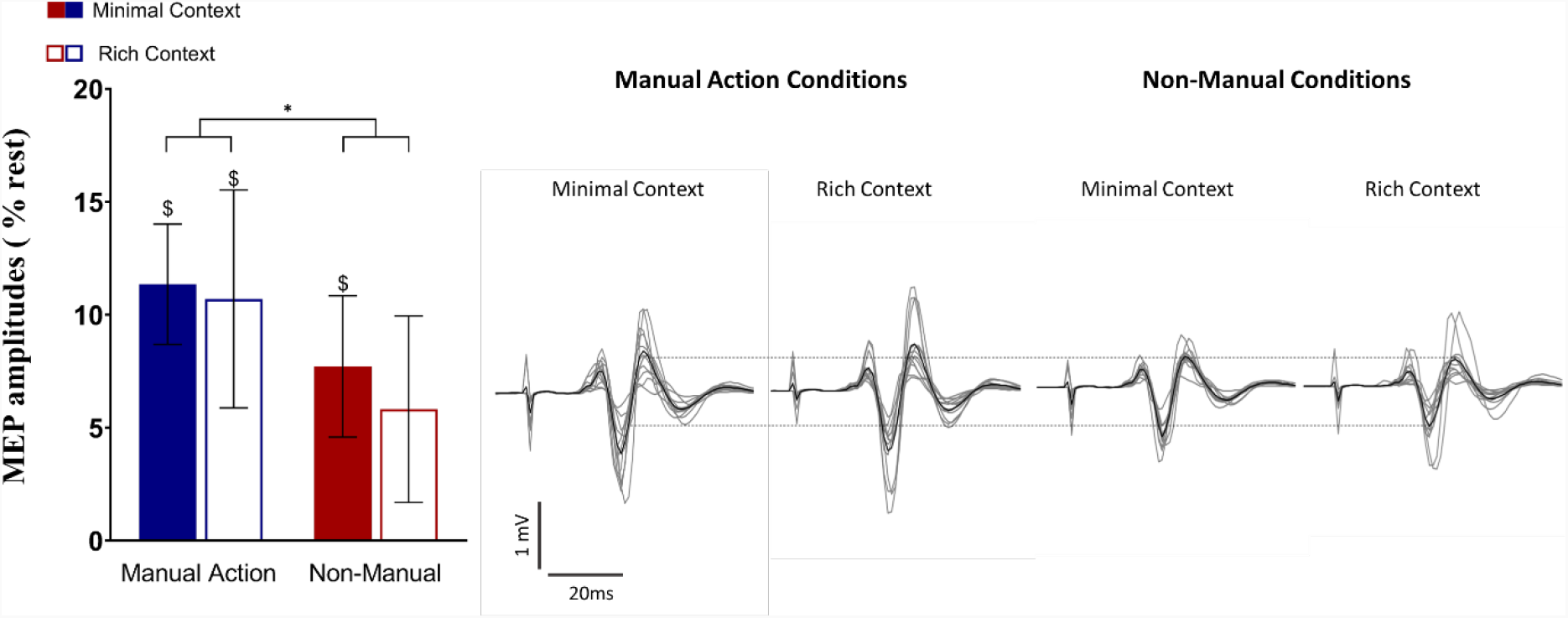
Corticospinal excitability in Rich and Minimal Context. Bar plots on the left side represent normalized MEPs (Vertical bars denote the Standard Error of the mean). Anova revealed an action effect for rich context but no effect for minimal context. The right side of the panel illustrates raw MEPs of a typical subject (grey lines). The black line is the average MEP of the condition for this participant. * = p<0.05. $=p<0.05 indicates a significant difference from zero (Trick trials).

Using one-sample t-tests, we found that normalized MEPs of all conditions were different from baseline (all p’s<0.05), except for non-manual verbs in the rich context (p=0.167). This finding reveals the importance of context for eliminating any residual activation for non-manual verbs that may be unclear in the absence of sentence context.

Low and high-level reading comprehension abilities influenced these outcomes, and in particular the context effect. We observed a positive correlation between the modulations of corticospinal excitability in rich context conditions and the low-level reading comprehension scores (p=0.048; r=0.341). Specifically, higher scores in memory-based comprehension yielded greater corticospinal excitability in rich context conditions (but not minimal, p=0.495; r=0.121). This finding reveals that the modulation of corticospinal excitability varies across individuals, and that comprehension ability may explain in part such variability. No significant correlations were observed for the high-level context-based comprehension scores (minimal context: p=0.623; r=-0.087; rich context: p=0.590; r=0.095) (Figure 3).

**Figure 3.**
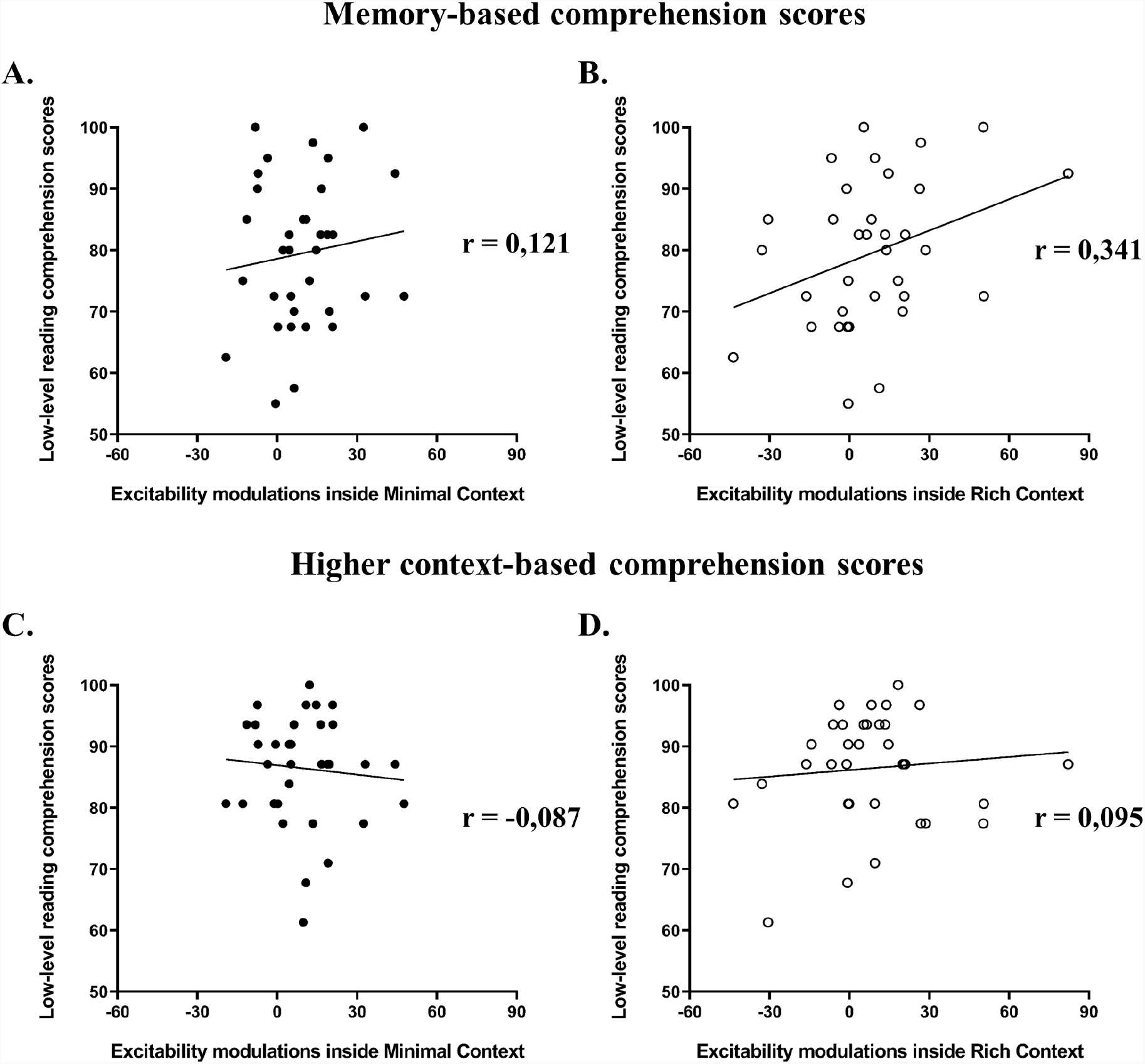
Correlations between the corticospinal excitability modulations depend on the context (Minimal vs. Rich) and the comprehension scores (memory vs. context-level). (A and B) represent correlations between the excitability modulations in Minimal context (A) and Rich context (B) with the memory-based comprehension scores. (C and D) represent correlations between the excitability modulations in Minimal context (C) and Rich context (D) with the higher context-level comprehension scores.

Finally, we analyzed the EMGrms prior to TMS artifacts to test for confounds in MEP modulation. We did not observe any difference between baseline (mean ±SD = 1.76 ±1.06µV) and the other conditions (all p’s>0.05; in minimal context, for manual verbs: 1.83 ±1.10µV, for non-manual verbs: 1.84 ±1.13µV; in rich context, for manual verbs: 1.90 ±1.48µV, for non-manual verbs: 1.83 ±1.11µV). This indicates that MEP modulations were not influenced by background EMGrms.

## 4. Discussion

The results of this study support the importance of the type of stimuli in modulating the involvement of the motor cortex during action language processing. We observed action-specific modulation of corticospinal excitability, with increased MEP amplitude for manual action verbs compared to non-manual verbs. Nonetheless, we did not observe a main effect of context nor an interaction between action and context. These findings do not support the “strengthening” idea, whereby sentence context would generate greater motor activation in general. However, there does seem to be some evidence for the idea of “sharpening,” as context was shown to minimize motor activation when sentence context makes it clear that there is no manual action.

The activation of the motor cortex is presumed to arise from reading events describing actions, typically verbs. However, even a non-action verb may be able to evoke some degree of motor activation if it is underspecified, because the event representation is vague. On the other hand, if the verb is well contextualized, the motor representation is more precise, yielding activation of only the appropriate effectors in the motor system that are relevant to the event. This is well reflected in our results where all conditions resulted in a higher state of corticospinal excitability from the baseline, except for non-manual verbs in the rich context. This suggests that the motor cortex can be automatically activated at some level even by non-action verbs in the absence of context [see also motor activation in abstract verbs (Sakreida et al., 2013) and metaphoric sentences (Desai et al., 2013; Reilly et al., 2019)]. While we did not observe any boost in motor activation when a rich context specifies the nature of the described situation, we did observe the elimination of any residual motor activation from non-manual verbs for which the representation may have been unclear in the absence of context. The rich context may lead to a disambiguation of the meaning of the verb, especially when the verb is vague or polysemous. Consistent with Mahon and colleagues (Mahon, 2015; Mahon & Caramazza, 2008), motor cortex activation may therefore play an important role in embellishing certain aspects of word meaning that are relevant in a given context.

Moreover, an action effect in minimal context is consistent with previous studies (Innocenti et al., 2014; Papeo et al., 2009), reporting that many manual action verbs would activate the motor cortex to a greater extent than non-manual verbs without context. However, we expected a greater difference for these same stimuli if accompanied by more contextual information to better specify the action, supporting the sharpening idea. Indeed, the manual nature of the meaning of some verbs is quite precise regardless of context (e.g., “I write”) whereas others can be rather vague (e.g., “I take”). As mentioned above, this vagueness can even introduce motor cortex activation in verbs that are intended to be non-action stimuli. Although we did not observe a greater effect of action in the rich compared minimal context, further research is currently underway to further test this idea of sharpening motor activation.

The present study suggests that the context effect is sensitive to individual factors, such as memory-based comprehension ability. We found that participants with higher comprehension ability exhibited a greater increase in excitability when reading rich-context sentences, regardless of verb type. This does not necessarily mean that context automatically strengthens motor activity (since no main effect of context was observed), but rather that the engagement of the motor cortex can vary across situations and across individuals, and may depend on parameters such as comprehension ability.

Based on the current findings and previous studies (Zwaan et al. 2010; Van Dam, Van Dijk, et al. 2012; Van Dam, van Dongen, et al. 2012; Gilead et al. 2013; Van Dam et al. 2014; Beauprez et al. 2018, 2020), language-induced motor representations are flexible and at least to some extent sensitive to linguistic context. It appears that context may play a role in strengthening motor activation for good comprehenders, as well as eliminating residual motor activation for non-action verbs. These findings are consistent with cognitive theories that accommodate semantic flexibility and top-down processes, whereby relevant features and alternative meanings are determined by the semantic context very early as each new word is encountered (e.g., Gentner 1981; Marslen-Wilson 1987; Tomasino and Rumiati 2013).

The question remains as to whether activation of the primary motor cortex is necessary to process action language (see also: Wurm and Caramazza, 2019). Researchers observed that disturbances of motor areas caused by repeated TMS during language processing can yield a decrease in performance, implying the engagement of the motor cortex in this process (Courson et al., 2018; Repetto et al., 2013; Vukovic et al., 2017; Willems et al., 2011). While understanding might be disturbed, diminished, or less effective, it is not completely blocked. Although language comprehension may be able to proceed without the involvement of the primary motor cortex, our findings suggest that it may play a role in the optimization of this process, refining linguistic understanding and making it more effective when necessary.

One potential limitation of the current study is the number of experimental trials per participant. Since we cannot know exactly when the motor representation will engage and reach its maximum for each subject, we have to probe at several latencies on separate trials, and select the peak afterwards. This can be considered a strength of our study in that we consider individual differences in processing time, but it can also be a limitation in that we reduce the number of experimental trials. In future investigations, we plan to overcome this limitation by calibrating each subject’s individual processing time prior to the test session, as well as employing continuous measures, such as EEG. Finally, while we strove to maximize the control between the rich and minimal context conditions, we could not avoid the additional pronoun which sometimes preceded the verb in the rich context condition. When this occurred, the target verb was rendered grammatically transitive in the rich context (*Je le frotte*; I scrub it) while it remained intransitive in the minimal context (*Je frotte*; I scrub). Otherwise, the probe screens were identical for the two conditions.

In conclusion, this study provides relevant information about the role of sentential context on primary motor cortex activation. We found that linguistic context serves to focus the motor representation of the described action, eliminating residual motor activation for non-action verbs. Moreover, the context effect appears to be sensitive to individual differences in reading comprehension. Consistent with the idea of semantic flexibility and top-down processes, when sentence context makes it clear that no manual action is evoked, the motor cortex does not contribute to the meaning representation.

## Author Contributions

Experiment design: WD, CML, FL, CP

Data collection: WD

Statistical analysis: WD, CML, FL

Manuscript preparation: WD, CP, CML, FL

## Data availability statement

All data from this study are available at https://osf.io/5zbmq/?view_only=82c93af470fd42be8ca33969b14f1c37

## Acknowledgement

The authors are very grateful to Cyril Sirandré for help with the custom Neurostim software and the experimental setup.

